# Comparative Morphometric Analysis Of Captive Jaguars Vs. Wild Jaguars (*Panthera onca*) In Venezuela

**DOI:** 10.1101/2021.05.04.442612

**Authors:** Israel Cañizales

## Abstract

In Venezuela the oldest, although anecdotical, record known of a captive jaguar is a male in the city of Maracay between 1918 and 1935. Body measurements were made on 22 jaguars kept in Venezuelan zoos between 1996 and 2009 to provide data on external morphometrics and compare with the measurements of 25 free-living animals, published by Hoogesteijn and Mondolfi (1992) and determine whether there are significant differences in the morphology between sexes of captive and free-living animals. The evaluation of morphometric variation was focused on three body measurements and body weight: Head-and-body length (HBL) and Tail length (Tl). Total body length (TBL) was calculated by adding HBL and Tl. Body weight (BW) was determined using a clock scale to the nearest gramme/kilogram. To determine the values all animals were anesthetized. The sexes were analyzed separately. All data were used for the calculation of descriptive statistics. Student t-tests were used with a significance level α 0.05 to establish if there is a significant difference between the measurements and weights of the sexes. The TBL and HBL in captive males were observed to be 8.78% and 12.13%, respectively, less than that of free-living males. A principal components analysis made it possible to assess the variability between HBL, TBL, and BW between sexes and between free-living/wild and captive animals. This study showed no significant differences in the BW of captives and wild females. Some captive males’ jaguars in this study may be thinner than normal, because of their diet and others may be fatter because of the lack of physical activity imposed by captivity. Body mass index (BMI) was used to contrast the difference in the average mass of each sex in both captive and free-living animals. Simple linear regressions and correlation coefficients were used to determine the relationship between body mass and male/female size. According to correlation coefficients, the change in weight that can be attributed to the ratio of weight to body length is 8.55 % for males and 9.83 % for females.

## INTRODUCTION

The jaguar *Panthera onca* (Linnaeus, 1758) is the largest felid in the Americas, and is the third largest of the genus *Panthera* Oken 1816, after the tiger (*P. tigris*) and the lion (*P. leo*). In Venezuela this species is listed in the Red Book of Venezuelan Fauna as Vulnerable (Jedrzejewski et. al., 2015). In general, jaguar weights range from 31 to 158 kg and their total body length from 154 to 241 cm (Emmons, 1997). In Venezuela, Hoogesteijn and Mondolfi (1992) recorded males with a mean weight of 96 ± 18 kg (range: 68 – 121 kg), and a mean length (including tail) of 212 ± 18 cm (range: 181 – 234 cm), and females with a mean weight of 56 ± 7 kg (range: 43 – 65 kg) and a mean length of 186 ± 7 cm (range: 176 – 196 cm). Linares (1998) recorded similar values to those of Emmons (1997) for weight and length.

Zoos around the world have held jaguars for more than a century. The oldest known record is a female at the Philadelphia Zoo in 1875 (Johnson, 2013; McMillan, 1995). By 1999 the International Species Information System - ISIS registered 239 individuals in 111 zoos around the world, of which 86.20% were born in captivity. Johnson (2013) registered in the Jaguar North American Regional Studbook 174 individuals in 71 zoos. Camelo (2014) registered in the National Studbook of Jaguar 24 individuals in 8 Colombians zoos. Recently, Jiménez González et. al. (2020) registre 38 jaguars in 8 Colombian zoos.

In Venezuela, the first record of a jaguar kept in a zoo dates from January 1946 (Johnson, 2013; McMillan, 1996) and was a female at El Pinar Zoo in Caracas. However, anecdotal reports indicate that a male was taken to the private zoo of then-Venezuelan President Juan Vicente Gómez in the city of Maracay between 1918 and 1935. Boher and Trebbau (1992) reported a total of 43 specimens (22 males and 21 females) in the zoological collections of Venezuela, indicating that in many of these establishment’s reproduction was relatively easy and even produced surpluses. The last available inventory of the number of jaguars kept in captivity in Venezuela is from 1998 reported by the National Foundation of Zoological Parks and Aquariums (FUNPZA), where a total of 35 animals (16 males and 19 females) are registered.

The exhibits enclosures of jaguars in Venezuelan zoos are limited in size and are less spatially complex than those environments experienced by animals in the wild. Noise, lighting, temperature and humidity, containment surface and substrate types, as well as exposure and resting areas, are all potential stressors of captive jaguars. In response to these stimuli, adrenocorticotrophic hormone (ACTH) is released, which acts directly on the adrenal cortex by stimulating the production of glucocorticoids (GC) and androgens. GC has a high activity in felids, which at high levels may alter behavior (anxiogenic effect, associated with aggressiveness or depression), including excessive grooming, hyperactivity, bulimia or suppression of appetite and suppression of the immune system, which favors the appearance of autoimmune syndromes, such as rheumatoid arthritis (Koscinczuk, 2014).

In addition to these environmental conditions, animals face a variety of changes but the most pronounced are associated with diet and nutrition. Many zoos feed jaguars a meat diet, supplemented with vitamins, without organs, skin, or connective tissue, unlike the diets of wild jaguars. Changes in diet can generate morphological variations in both shape and dimensions. Studies of pantherine felids suggest that differences in the shape of the skull between captive and free-living individuals are related to differences in chewing load, resulting from differences in diet (Hartstone-Rose et. al., 2014; Zuccarelli, 2004). An abnormal ratio of calcium: phosphorus in the diet may be responsible.

Although morphological differences between wild and captive individuals have been observed in different mammal species over the years, these differences, and the impact of captivity on the anatomy of animals rarely have been studied (Hartstone-Rose et. al., 2014; O’Regan and Kitchener, 2005). Some studies conducted to assess the degree of difference have been focused mainly on primates (Altmann et al., 1993; Bolter and Zihlman, 2003; Lewton, 2017; Phillips-Conroy and Jolly, 1988; Turner et al., 2016), carnivores (Zuccarelli, 2004; Hailemariam et al., 2015; Saragusty, 2014; Weber Rosas et al., 2009) and mice (Courtney Jones et al., 2018; McPhee, 2004).

One of the physical changes that can result in generations of captive-born animals is a reduction in sexual dimorphism, which in turn is related to the rate or age of maturity. Alternatively, morphological differences may occur due to changes in the strength and targets of selection in captivity, selecting morphological phenotypes that maximize fitness in captivity (Mathews et al., 2005; Schulte-Hostedde and Mastromonaco, 2015). This cannot be interpreted as “adaptive” in an evolutionary sense, since there is not enough time for selective pressures to act, although it could well be “adaptive” in terms of a phenotypic response to a different environment, but not fixed genetically.

On the other hand, obtaining accurate data on external measurements and weight in live jaguars both in captivity and in the field is perhaps the main difficulty to overcome. The difficulty in handling and obtaining reliable data in jaguars will depend on the immobilization techniques and the experience of the recorder to reduce the risk of accidents and measurement errors. For some researchers the skins deposited in museums should not be measured. Because skins can be affected by the processes of preparation and fixation of the tissues, with some influence of the statistical analyses commonly used.

Based on the above, this study provides biometric data of captive jaguars collected between 1996 and 2009 and its comparison with the data published of free-living ones by other authors to determine if there is significant variation in external morphometry and body mass between captive and wild animals in Venezuela.

## MATERIALS AND METHODS

The morphometric analysis was made based on 22 jaguars held in captivity in 10 facilities (nine zoos and one private collection) (Fig. 1). All the animals were adults, between three and 17 years old. For the comparison of body measurements with the captive animals were used the data published by Hoogesteijn and Mondolfi (1992) from 25 free-living Venezuelan jaguars.

**Figure 1.**
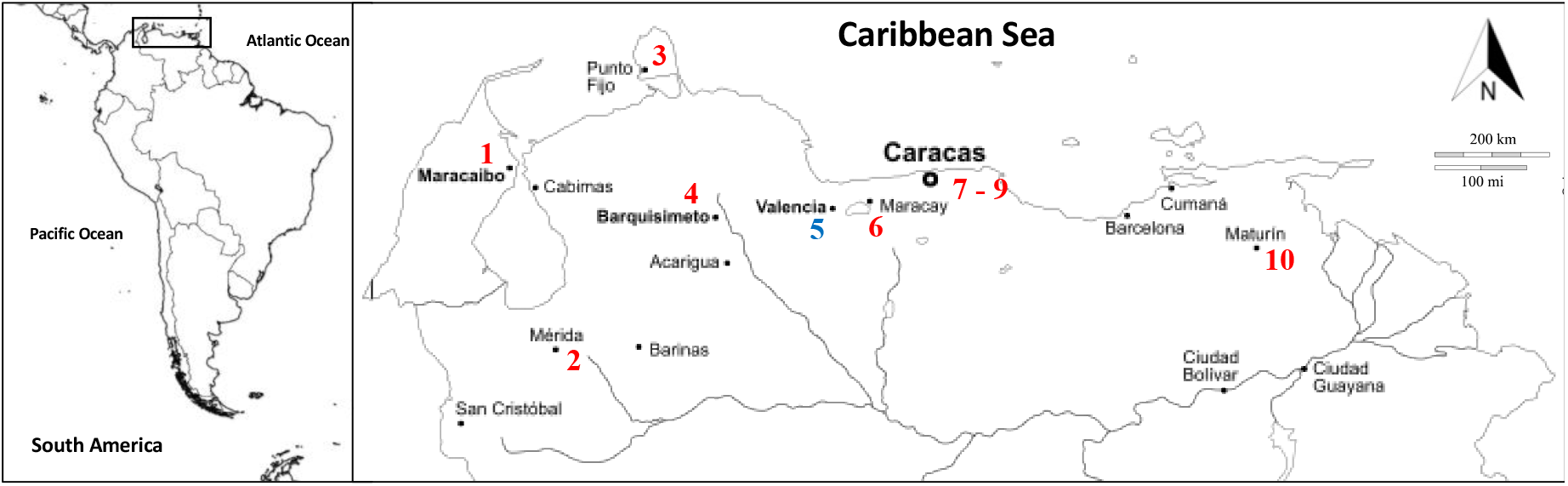
Relative geographic location of sampling localities. Red = Zoos, Blue = Private collection. 1 = Metropolitano del Zulia, 2 = Chorros de Milla, 3 = Paraguaná, 4 = Bararida, 5 = Association of Cattle Breeders of Valencia, 6 = Las Delicias, 7 = Caricuao, 8 = El Pinar, 9 = Generalísimo Francisco de Miranda, 10 = La Guaricha.

*Linear morphometry.* The evaluation of morphometric variation in this study focused on three body measurements and body weight. The point-to-point measurements were: Head-and-body length (HBL) = from the tip of the muzzle/nose to the base of the tail, and Tail length (Tl) = from the base of the tail to the end of the last caudal vertebra, excluding the terminal tuft of hair. Total body length (TBL) was calculated by adding HBL and Tl (Fig. 2). The measurements were taken by the author with the anesthetized animals following an anesthetic protocol with the use of a xylazine-ketamine combination as described by Cañizales (2019) and using tape measures calibrated in centimeters (cm) with two or three repetitions to an accuracy level of ±1.0 cm to minimize recorder error. Body weight (BW) was recorded using a clock scale, which can weigh up to 200 kg.

**Figure 2.**
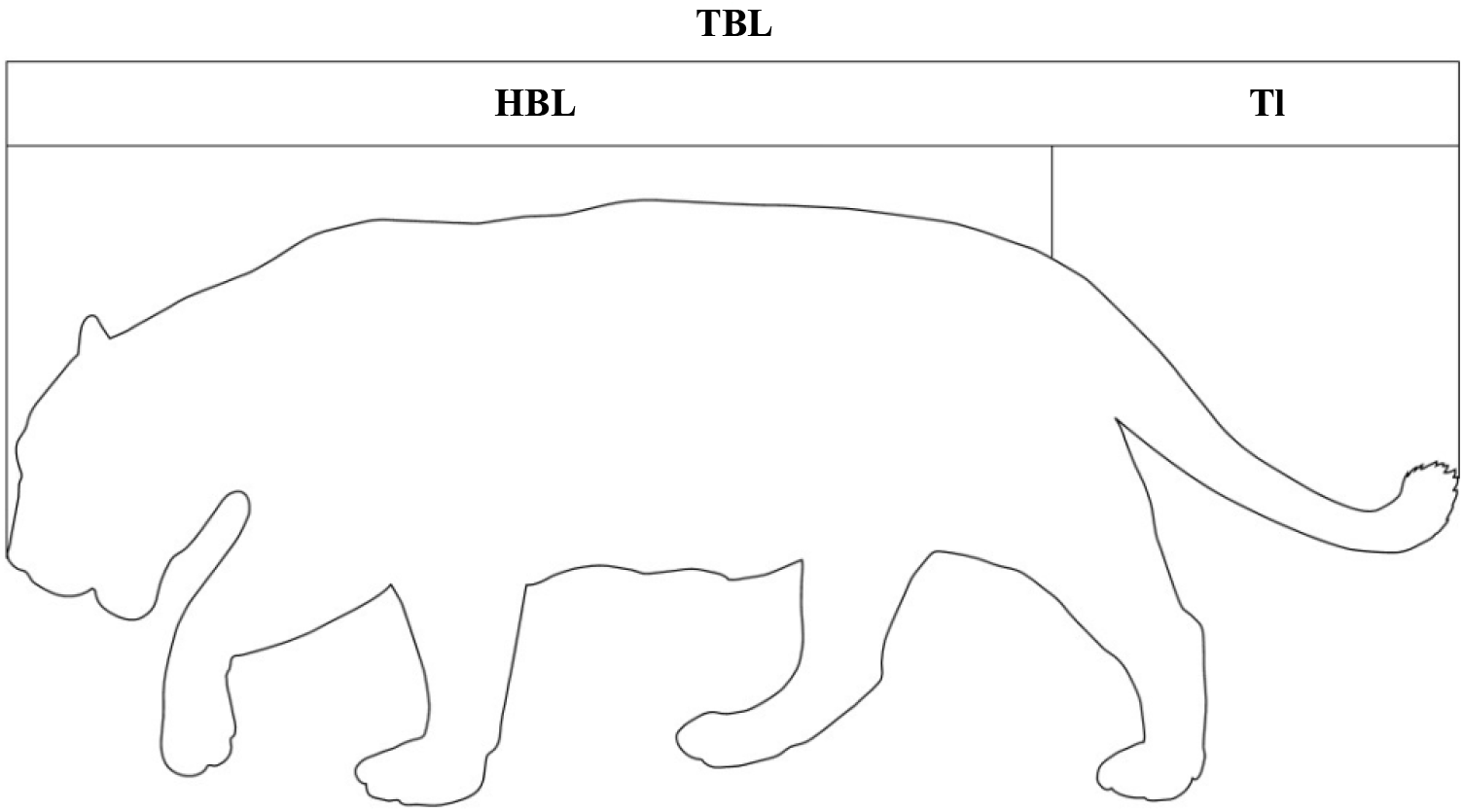
Schematic drawing showing external body measurement points in jaguars. (TBL) total body length, (HBL) head-body length, (Tl) tail length.

All data were used in calculations of descriptive statistics (Mean, Standard Deviation, Minima and Maxima, Coefficient of Variation). The sexes were analyzed separately. The *t-* Student test (p ≤ 0.05) was used to determine if there are significant differences in morphometric data between males and females both in captivity and in the wild, as well as between animals of the same sex in both situations. Violin plots were generated to illustrate size differences. In addition, and as an exploratory descriptive tool, a principal component analysis (PCA) was performed. This multivariate technique considers different variables to determine the patterns of morphometric variation between groups, as well as to evaluate the degree of separation between them, trying to achieve maximum homogeneity so that the forms are grouped according to the degree of similarity. The graphs were obtained using the PAST 4.03 program (Hammer et. al., 2001).

*Body condition.* The following scoring systems were used to assess and assign the degree of body symmetry and muscle development: Body condition score (BCS) which evaluates body fat coverage by visual estimation on a 1–9 scale (1 = Very thin, No detectable body fat, and 9 = Obese, Heavy fat cover), and Muscle condition score (MCS), which evaluates by touch on a 1–4 scale the firmness or turgidity of the muscle masses around the temporal bones, scapulae, lumbar vertebrae, and pelvic bones (1 = No muscle wasting, normal muscle mass, and 4 = Marked muscle wasting). Both scores were based on the criteria of AZA (2016), Baldwin et. al. (2010), and Laflamme (1997). To contrast the body mass ratio between captive and free-living animals, a Body mass index (BMI) was calculated. For this purpose, was used the formula:

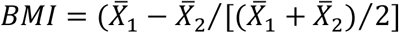

where 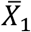 is the mean weight in sample 1, and 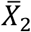 is the mean weight in sample 2. This formula provides the difference in mass relative to the average mass of the individuals (De La Torre and Rivero, 2017). To model possible differences between males and females, body weight and the lengths measures were analyzed using linear regression.

## RESULTS

A total of 47 jaguars of known captive or wild status were included in this study. These comprise 27 males and 20 females. The males consisted of 12 captive animals and 15 of free-living ones whilst for females there were 10 of captive and wild origin, respectively.

*Linear morphometry.* Table 1 shows all measurements in centimeters and weights in kilograms obtained in this study for captive animals and those reported by Hoogesteijn and Mondolfi (1992) for free-living animals in Venezuela. The animal with the largest TBL was a free-living male with 234 cm. The HBL, TBL and BW of captive males show mean values lower than those of free-living males. In females the HBL variable presents a slightly higher mean value in captive animals. No significant differences were found in CV between captive and wild jaguars [Males: *t =* −0.0910, *DF =* 6, *p =* 0.4652, captivity (3 values ≥ 10%); free-living ones (2 values ≥ 10%)]; [Females: *t =* 0.9769, *DF =* 5, *p =* 0.1867, captivity (2 values ≥ 10%); free-living ones (1 value ≥ 10%)]. All variables between captives and free-living males were shows significant differences (TBL: *t =* 2.451; *DF =* 22; *p =* 0.011; HBL: *t =* 2.590; *DF =* 17; *p =* 0.010; BW: *t =* 5.423; *DF =* 24; *p =* 0.000). In the case of females all variables show no significant differences (TBL: *t =* 1.681; *DF =* 16; *p =* 0.056; HBL: *t =* −0.209; *DF =* 17; *p =* 0.418; BW: *t =* 0.947; *DF =* 16; *p =* 0.179).

**Table 1.** Mean and standard deviation of body measurements and body mass of captive and free-living jaguars discriminated by sex. All measurements are in centimeters. Body mass is in kg.

| Variable | Captive jaguars (CJ) |  |  |  |  |  | Free-living jaguars (FLJ) |  |  |  |  |  |
| --- | --- | --- | --- | --- | --- | --- | --- | --- | --- | --- | --- | --- |
|  | This study |  |  |  |  |  | Hoogesteijn and Mondolfi (1992) |  |  |  |  |  |
|  | Males |  |  | Females |  |  | Males |  |  | Females |  |  |
| | Mean ± SD | Min – Max (*n*) | CV | Mean ± SD | Min – Max (*n*) | CV | Mean ± SD | Min – Max (*n*) | CV | Mean ± SD | Min – Max (*n*) | CV |
| TBL | 193.08 ± 20.51 | 138 – 216 (12) | 0.106 | 179.40 ± 9.56 | 161 – 196 (10) | **0.053** | 211.67 ± 18.34 | 181 – 234 (15) | **0.087** | 185.60 ± 6.69 | 176 – 196 (10) | **0.036** |
| HBL | 131.25 ± 21.19 | 70 – 152 (12) | 0.161 | 127.30 ± 7.18 | 115 – 138 (10) | **0.056** | 149.36 ± 12.68 | 126 – 170 (15) | **0.085** | 126.70 ± 5.54 | 116 – 134 (10) | **0.044** |
| Tl | 61.83 ± 4.11 | 56 – 70 (12) | **0.066** | 52.10 ± 7.75 | 41 – 62 (10) | 0.149 | 60.79 ± 9.89 | 33 – 71 (15) | 0.163 | 58.90 ± 5.13 | 50 – 66 (10) | **0.087** |
| BW | 65.50 ± 11.33 | 45 – 87 (12) | 0.173 | 52.30 ± 9.56 | 40 – 70 (10) | 0.183 | 95.87 ± 17.61 | 68 – 121 (15) | 0.184 | 55.89 ± 6.86 | 43 – 65 (9) | 0.123 |

TBL = Total body length, HBL = Head-body length, Tl = Tail length, BW = Body mass, SD = Standard deviation, Min = Minima, Max =Maxima, (*n*) = sample size, CV = coefficient of variation. (**Bold** = lower variation).

For captive males the mean TBL was significantly longer than that of females (*t =* 2.058; *DF =* 16; *p =* 0.028). The mean weight of the males was significantly greater than that of the females (*t =* −2.965; *DF =* 20; *p =* 0.004). However, the mean HBL of captive males was not significantly longer than that of captive females (*t =* −0.605; *DF =* 14; *p =* 0.277). In free-living animals all variables in males were significantly greater than those of females (TBL: *t =* 5.305; *DF =* 18; *p =* 0.000; HBL: *t =* 6.092; *DF =* 19; *p =* 0.000; BW: *t =* 7.835; *DF =* 20; *p =* 0.000).

The violin diagram allows to visualize and demonstrate the differences in size and the multimodal distribution of values by sex both in captivity and in the wild (Fig. 3). The mean TBL and mean HBL of captive males were significantly less than those of free-living males (*t* = 2.451; DF = 22; *p =* 0.011; *t =* 2.590; *DF =* 17; *p =* 0.010). However, the mean TBL and mean HBL of captive females are not significantly different from those of free-living females (*t =* 1.681; *DF =* 16; *p =* 0.056; *t =* −0.209; *DF =* 17; *p =* 0.418), respectively.

**Figure 3.**
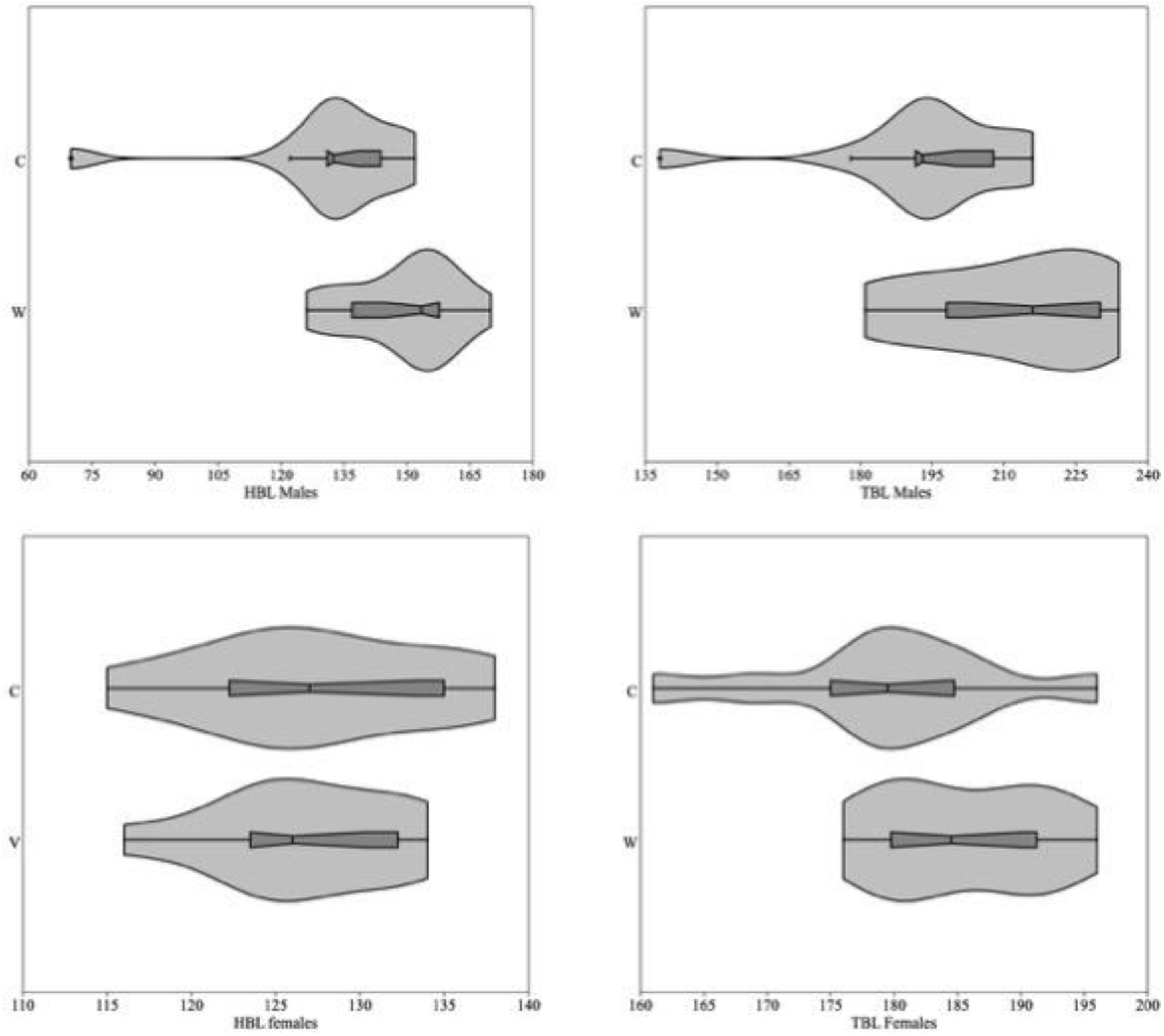
Violin plot of HBL and TBL of captive and free-living jaguars. The central thick black bar represents the interquartile range. The left and right thin black lines extending from it represent the 95 % confidence intervals. The gray areas represent the distribution of values.

In the PCA the percentage of the total variation that best explains the results is concentrated in the first two components (96.93%). The plot shows overlap between captive and wild specimens (Fig. 4). PC1 explains 83.25% of the variance by grouping females and most captive males with free-living females by total body weight and length. While free-living males with some captive males remain on the positive side of the axis, which means that these are much larger in these characteristics. PC2 explains 13.67% of the variance and groups individuals by body length (Fig. 4).

**Figure 4.**
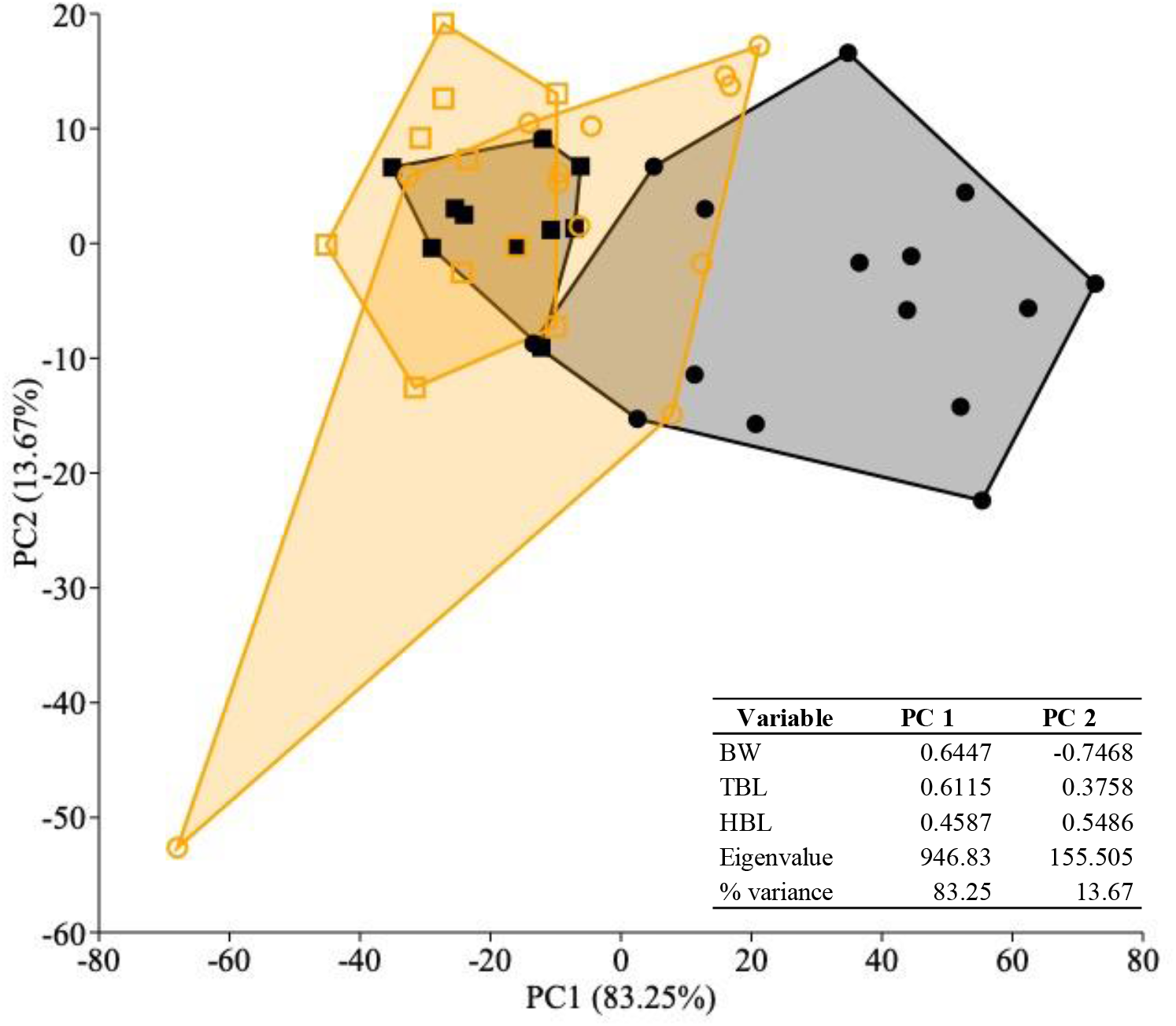
Bivariate plot of principal component analysis. (•) Free-living males, (□) Free-living females, (Ο) Captive males, (□)females in captivity. Lines were drawn around each group to aid visualization. The loadings, eigenvalues and % variance of the variables are shown.

*Body condition.* The maximum weight recorded for captive jaguars in this study was 87 kg for a male and 70 kg for a female. In free-living jaguars, the maximum weight recorded was 121 kg for a male and 65 kg for a female (Hoogesteijn and Mondolfi, 1992). The BW of females in captivity does not differ significantly from that of wild females (*t =* 0.947; *DF =* 16; *p =* 0.179), unlike captive males (*t =* 5.423; *DF =* 24; *p =* 0.000), which were significantly less heavy than wild males.

The 22 captive jaguars were assigned BCS ranging from 3 to 5 and MCS ranging from 2 to 3. Females and males were equally distributed in all BCS categories except for as light but significant overrepresentation of males with a BCS of 5, compared with females with the same score (Table 2). The BMI of captive versus free-living jaguars by sex indicates that free-living males were 0.38 times heavier. Similarly, the BMI of free-living females indicates that they are slightly heavier with a value of 0.07.

**Table 2.**
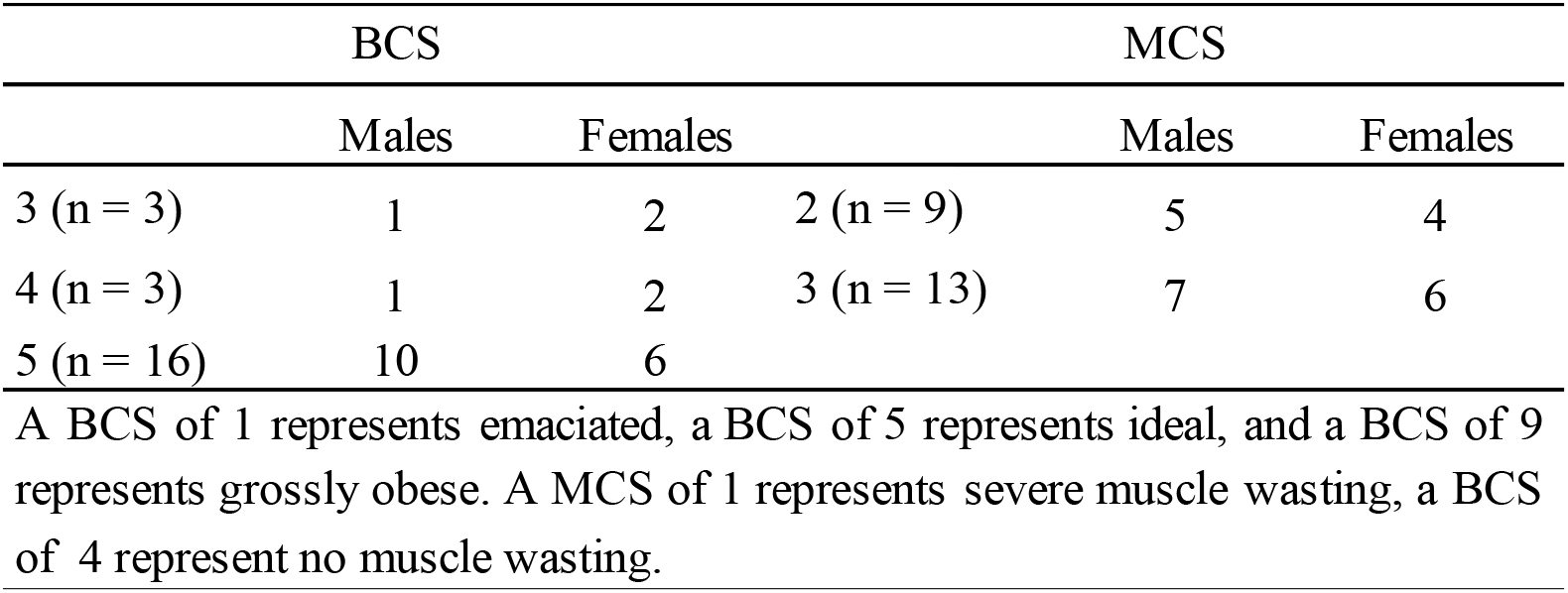
BCS and MCS of captive discriminated by sex assigned based on visual and palpatory evaluation.

Figure 5 (a and b) shows the relationship and linear regression lines between total body length (TBL) and body weight (BW) of males and females, respectively. In both cases, a moderate heterogeneous distribution is observed, although with a certain linear trend and a positive upward slope. Although some points are moderately off the line, the mean values of males’ jaguars in captivity (red dots) are lower than those free-living (black dots).

**Figure 5.**
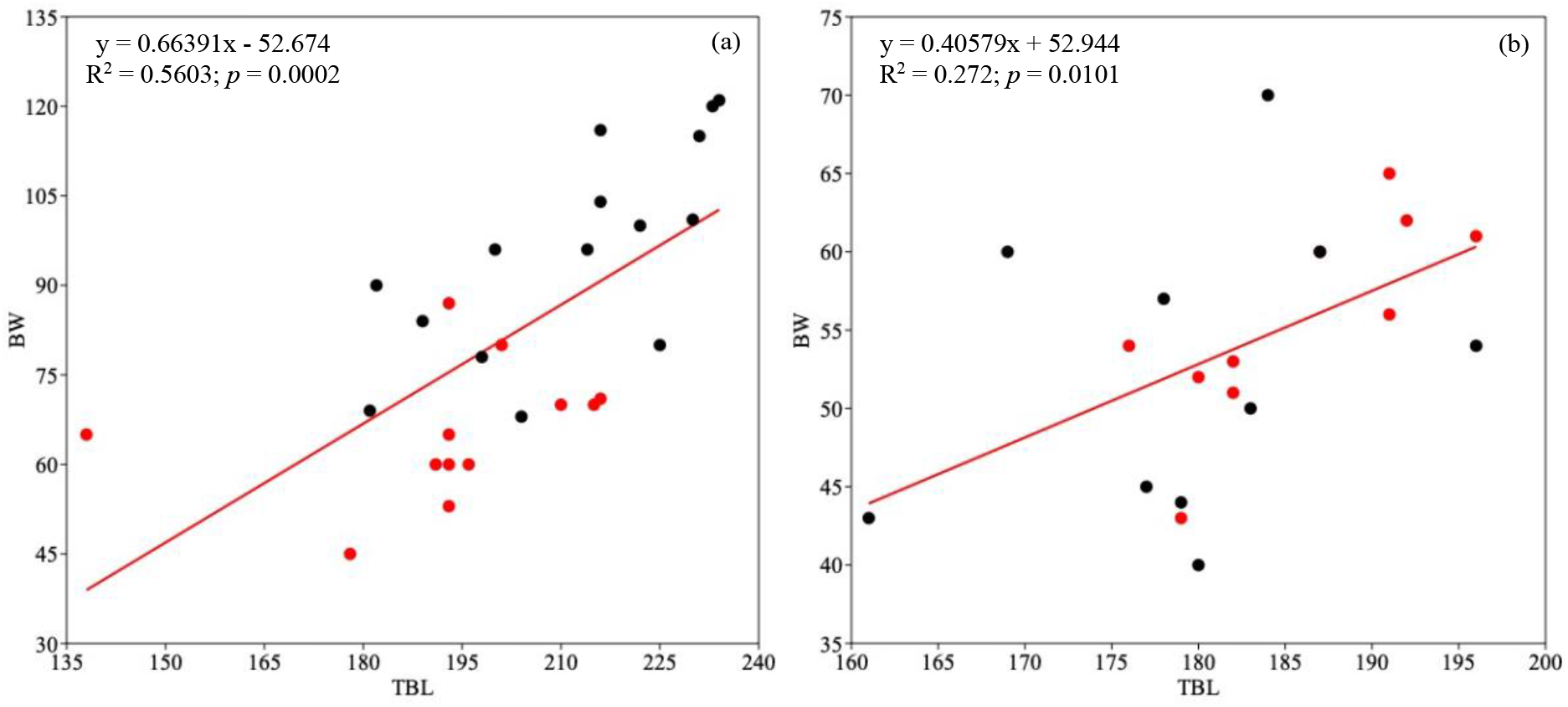
Linear regression between BW and TBL of male (a) and female (b) jaguars. The red dots represent captive animals. The black dots represent free-living animals. The values of slope, intercept and correlation coefficients are reported.

Correlation coefficients for each sex are 0.5603 for males and 0.272 for females, in other words 56.03% of the variation in body weight in males that can be attributed to the relationship with total body length, indicating that the relationship between these variables is considered moderate to strong. On the other hand, in females only 27.2% of the variation in weight can be attributed to the relationship with total body length, indicating that the relationship between these variables is considered weak.

## DISCUSSION

For many endangered mammal species, captive animals are the only individuals available for reproductive, behavioral, feeding, and other studies, including their morphology. And although in some cases the number of live animals may not always satisfy the statistical assumptions, they can explain the biological assumptions under study. On the other hand, the capture or chemical immobilization in the field of elusive species such as the jaguar requires time, properly trained personnel, equipment, logistics and economic resources that in most cases are prohibitive. Additionally, the number and quality of samples (i.e., skins) in museums and collections are variable, and it would be illegal and unethical to collect new biological material to meet the statistical criteria for validating taxonomic or morphological assessments.

In this situation, the question arises: what is the alternative? Either do not study any species, because there is insufficient material (i.e., captive, or wild) available or try to understand what structures may have been altered by captivity and quantify that change. Unfortunately, the prevailing opinion is often the first.

Despite the above, morphological differences between free-living and captive adult animals have been reported in several mammalian species (Courtney Jones et. al., 2018; O’Regan and Kitchener, 2005; Turner et. al., 2016).

*Linear morphometry.* In the present study, the mean TBL and HBL in captive males were observed to be 8.78% and 12.13%, respectively, lower compared to free-living jaguars recorded by Hoogesteijn and Mondolfi (1992). While among females, no significant differences were observed.

The differences found may be due to biological (intrinsic rate of growth related to age) and environmental factors (housing conditions, changes in diet), including measurement errors or simply be due to population differences. In most terrestrial mammals there are some external criteria for determining age, such as type and color of fur, eruption and type of teeth, and primary and secondary sexual characteristics, as appropriate. However, there are other biological elements that can be used in captivity, such as the evaluation of the opacity of the crystalline lens, the ossification of the ossicles of the ear, the evaluation of the corpora lutea in females, as well as diagnoses based on radiographic techniques. This technique evaluates the ossification points of both long and short bones. Whether of cartilaginous or membranous origin, it is the appropriate intake of protein, calcium, and phosphorus, the hormonal balance, and the state of health that will determine the final degree of ossification and the final body lengths (Zoran, 2002). Venezuelan zoos usually feed jaguars with meat (beef, chicken, horse, or pork), and in some cases supplemented with mineral or vitamins, without organs, unlike the diets of wild jaguars, with a feeding schedule every two days.

The maximum TBL recorded by Hoogesteijn and Mondolfi (1992) was in a male with 234 cm. In contrast, Emmons (1997) for wild jaguars in Neotropic register a TBL 241 cm and Linares (1998) referred the same TBL for wild jaguars in Venezuela, however these measurements were obtained from skins and by different observers without a precise description of how they were measured. During the preservation or tanning process of skins distortions and alterations can occur, and probably the measurements are not representative it all. Moreover, measurements taken from skins of animals hunted from the end of the 19th to the middle of the 20th centuries, may vary considerably. In this period, most measurements followed the curvilinear contour of the body and these curvilinear lengths are always longer than those measured in a straight line (standard length). However, this does not seem to be the case for females whose lengths are like the measures reported for females in the wild in Venezuela by Hoogesteijn and Mondolfi (1992).

Considering that the free-living males jaguars come from different regions of Venezuela and significant differences were found, it can be concluded that the maximum TBL and HBL reached by the captive males in this study can be partially explained by the effect of captivity, as suggested by O’Regan and Kitchener (2005).

*Body condition.* In this study, the mean BW of captive males (65.50 kg) was 31.68% lower than the mean BW of free-living ones recorded by Hoogesteijn and Mondolfi (1992). This could be considered unexpected because captives animals tend to be much fatter (and less muscular) than their wild counterparts. Although, this is more often observable in developed countries where the food supply is guaranteed all the year round, should be considered the fact that in Venezuelan zoos the amount of food is generally deducted based on 3.0 % of the live weight of the animal delivered in a every other day food ration. However, the mean BW of females were 6.42% heavier than their free-living counterparts. In this case some females may be fatter due to the limitation of physical activity imposed by captivity. In addition, larger females may be better able to protect their offspring and deterring infanticide.

Although the results of this study show no significant differences in the BW of captive and wild females, it is possible to state that the relationship between TBL and BW, owing principally to sample size, particularly in females, not increase in a statistically significant way. It is also possible that these differences are due to variations in the methods of weighing.

A limitation of both BCS and MCS systems was the risk of investigator bias. The BCS and the MCS are not directly related, an animal may be overweight, but may have significant muscle loss. This can make a low to moderate value MCS look relatively normal, if not carefully evaluated. Although BCS is a subjective evaluation based on clinical examination but despite its subjective basis, a good correlation of scores reportedly exists between assessors (AZA, 2016; Baldwin et al., 2010; Laflamme, 1997).

In these cases, although some areas of the body may appear relatively normal or even fat-rich (especially on the ribs or in the abdominal region), muscular atrophy makes bone structures evident. Considering the above, the BCS associated with the MCS the captive animals placed 16 in category 5 and 13 in category 3 respectively and since they do not have reference values, it is assumed that the value obtained in this study is probably the maximum reached by the species in captivity in Venezuela.

In this study the BMI calculated between captive males and females with free-living jaguars’ counterparts were 0.38 and 0.07, respectively. If we calculate and compare the BMI for free-living animals in Venezuela according to data published by Hoogesteijn and Mondolfi (1992), gives a result of 0.53, slightly higher than the value published by De La Torre and Rivero (2017) of 0.42 for free-living jaguars in Mexico. The variation may be related to the availability of larger prey (Hoogesteijn and Mondolfi 1992).

The differences observed in the values associated with BMI between captive and free-living animals could be based on the availability of food resources and physical activity. In captivity activity levels differ markedly from those in nature. Although the variables of size and type of soil or substrate of enclosures were not evaluated in this work, they partly determine the degree of activity and consequently the muscular development or fat accumulation of the animals. Captive animals, on the other hand, are routinely supplied and do not need to spend energy searching for food.

The results here confirm the existence of differences of captive jaguars in comparison to those of wild. Although the sample in this study may be considered small, significant differences in the TBL, BL and BW between adult males jaguars captives and free-living was found. Whether these differences are related to insufficient nutrients or to other environmental factors in captivity and/or to genetic factors has yet to be determined.

## ACKNOWLEDGEMENTS

To Venezuelan zoos for allowing access to their facilities. To A. Blanco, D. Garcia, A. Henríquez, R. López, J.M. Pernalete, A. Quintero and M. Santana for their support and assistance in all these years of work. To the memory of my beloved son Armando.

## ETHICAL STATEMENT

To carry out the study, approval was obtained through an informed consent read and signed by the administrative representatives of each facility. All protocols followed good practices and animal welfare principles set forth in the Law on the Practice of Veterinary Medicine Official Gazette No. 28,737 dated 24 September 1968 and the Law on the Protection of Wild Fauna Official Gazette No. 28,289 dated 11 August 1970 and approved by Animal Research Ethical Committee. All efforts were made to utilize only the minimum number of animals necessary to produce reliable scientific data.

## CONFLICT OF INTEREST

The author declares that there is no conflict of interest.

## Notes

### Competing Interest Statement

The authors have declared no competing interest.

## REFERENCES

Altmann, J., D. Schoeller, S.A. Altmann, P. Muruthi and R.M. Sapolsky. 1993. Body size and fatness of free-living baboons reflect food availability and activity levels. Am J Primatol. 30:149–161.

AZA Jaguar Species Survival Plan. (2016). Jaguar Care Manual. Silver Spring, MD: Association of Zoos and Aquariums.

Baldwin, K., J. Bartges, T. Buffington, L.M. Freeman, M. Grabow, J. Legred and D. Ostwald. (2010). Guía para la evaluación nutricional de perros y gatos de la Asociación Americana Hospitalaria de Animales (AAHA). Journal of the American Animal Hospital Association. 46:285–297.

Boher, S., and P. Trebbau. (1992) El papel de los parques zoológicos modernos en la conservación de los Yaguares en Venezuela. In: Felinos de Venezuela. Biología, Ecología y Conservación. Fundación para el Desarrollo de las Ciencias Físicas, Matemáticas y Naturales. Caracas. Venezuela. p.301–305.

Bolter, D.R. and A.L. Zihlman. 2011. Brief communication: Dental development timing in captive Pan paniscus with comparison to Pan troglodytes. Am J Phys Anthrop. 145:647–652. [PubMed: 21541924]

Camelo, V. 2014. Studbook Nacional de Jaguar (Panthera onca). 23 pp.

Cañizales I. 2019. Inmovilización química, hematología y química sanguínea de jaguares (Panthera onca) en zoológicos de Venezuela: estudio retrospectivo, 1996-2009. Rev Med Vet. (38): 47–62. doi: https://doi.org/10.19052/mv.vol1.iss38.5

Courtney Jones, S.K., J. Munn Adam and P.G. Byrne. 2018. Effect of captivity on morphology: negligible changes in external morphology mask significant changes in internal morphology. R. Soc. open sci.5172470172470 http://doi.org/10.1098/rsos.172470

De La Torre, J.A. and M. Rivero. 2017. A morphological comparison of jaguars and pumas in southern Mexico. THERYA. 8:117–122.

Emmons, L. 1997. Neotropical Rainforest Mammals. A Field Guide 2nd Ed. The University of Chicago Press. Chicago. EE.UU.

Hailemariam, D., L. Alemayehu and T. Yilma. 2015 Reproductive Characteristics and Body Morphometry of Captive Lions (Panthera leo) at Addis Ababa Zoo. World Journal of Zoology. 10 (3): 226–232. DOI: 10.5829/idosi.wjz.2015.10.3.95102

Hammer, Ø., D.A.T. Harper and P.D. Ryan. 2001. PAST: Paleontological Statistic software package for education.

Hartstone-Rose, A., H. Selvey, J.R. Villari, M. Atwell and T. Schmidt. 2014. The three-dimensional morphological effects of captivity. PLoS ONE 9(11): e113437. doi: 10.1371/journal.pone.0113437

Hoogesteijn, R. and E. Mondolfi. (1992) El Jaguar. Armitano Publishers. Caracas. Venezuela. International Species Information System. 1999. Animal record keeping system. Apple Valley, Minnesota. EE. UU.

Jiménez, S., H. Monsalve, M.A. Moreno and C. Jiménez. 2020. Demographic analysis for the reproductive management of captive jaguars (Panthera onca) in Colombian zoos. Biota Colombiana. 21(1): 86–103. DOI: 10.21068/c2020.v21n01a06.

Johnson, S. 2013. AZA Regional studbook jaguar (Panthera onca). EE. UU

Koscinczuk, P. 2014. Ambiente, adaptación y estrés. Revista Veterinaria. 25:67–76.

Laflamme, D. 1997. Development and validation of a body condition score system for cats: a clinical tool. Feline Practice. 25:13–18.

Lewton, K.L. 2017. The effects of captive versus wild rearing environments on long bone articular surfaces in common chimpanzees (Pan troglodytes). PeerJ 5:e3668; DOI 10.7717/peerj.3668

Linares, O. 1998. Mamíferos de Venezuela. Sociedad Conservacionista Audubon de Venezuela. Caracas. Venezuela.

Mathews, F., M. Orros, G. McLaren, M. Gelling and R. Foster. 2005. Keeping fit on the ark: assessing the suitability of captive-bred animals for release. Biol. Conserv. 121, 569–577. (doi: 10.1016/j.biocon.2004.06.007).

McMillan, G. 1996. Jaguar north American regional studbook. USA

McPhee, M.E. 2004. Morphological change in wild and captive old field mice Peromyscus polionotus subgriseus. J. Mammal. 85, 1130–1137. (doi:10.1644/BPR-017.1)

O’Regan, H.J. and A.C Kitchener. 2005. The effects of captivity on the morphology of captive, domesticated and feral mammals. Mammal Review. 35:215–230.

Phillips-Conroy, J.E and C.J. Jolly. 1988. Dental eruption schedules of wild and captive baboons. Am J Primatol. 15:17–29.

Jedrzejewski, W., M.R. Abarca-Medina, E.O. Boede, R. Hoogesteijn, E. Isasi-Catalá, R. Carreño, A.L. Viloria, H. Cerda, D. Lew, A.J. González-Fernández, L. Perera and M.F. Puerto Carrillo. 2015. Jaguar, Panthera onca, In: J.P. Rodríguez, A. García-Rawlins and F. Rojas-Suárez (eds.) Libro Rojo de la Fauna Venezolana. Cuarta edición. Provita y Fundación Empresas Polar, Caracas, Venezuela., Recuperado de: www.especiesamenazadas.org/taxon/chordata/mammalia/carnivora/felidae/panthera/jaguar.

Saragusty, J., A. Shavit-Meyrav, N. Yamaguchi, R. Nadler, T. Bdolah-Abram et al. 2014. Comparative Skull Analysis Suggests Species-Specific Captivity-Related Malformation in Lions (Panthera leo). PLoS ONE 9(4): e94527. doi: 10.1371/journal.pone.0094527

Schulte-Hostedde, A.I. and G.F. Mastromonaco. 2015. Integrating evolution in the management of captive zoo populations. Evol. Appl. 8:413–422. (doi:10.1111/eva.12258)

Turner, T. R., Cramer, J. D., Nisbett, A. and Patrick Gray, J. 2016. A comparison of adult body size between captive and wild vervet monkeys (Chlorocebus aethiops sabaeus) on the island of St. Kitts. Primates. Journal of Primatology. 57(2): 211–220. https://doi.org/10.1007/s10329-015-0509-8

Weber Rosas, F.C., C. Soares da Rocha, de Mattos, G.E. and S.M. Lazzarini. 2009. Body weight-length relationships in giant otters (Pteronura brasiliensis) (Carnivora, Mustelidae). Brazilian Archives of Biology and Technology, 52(3), 587–591. https://doi.org/10.1590/S1516-89132009000300010

Zoran, D.L. 2002. The carnivore connection to nutrition in cats. Journal of American Veterinary Medical Association. 221:1559–1567.

Zuccarelli, D. 2004. Comparative morphometric analysis of captive vs. wild African lion (Panthera leo) skulls. Bios. 75:131–138.

